# Combined transcriptome and metabolome analysis of a new species of microalgae from Tibetan plateau and its response to sewage treatment

**DOI:** 10.1101/2022.06.16.496493

**Authors:** Wang Jinhu, Zhang Qiangying, Chen Junyu, Zhou Jinna, Li Jing, Wei Yanli, Bu Duo

**Affiliations:** College of Science, Tibet University, 10 East Zangda Road, Lhasa City, Tibet Autonomous Region, P.R. China 850000

**Keywords:** Qinghai-Tibet plateau, *Desmodesmus*, sewage treatment, transcriptomics, metabolomics, correlation analysis

## Abstract

Microalgae are pivotal in maintaining water quality in the lakes and rivers of Qinghai-Tibet plateau. The optimum sewage treatment conditions for *Desmodesmus* sp. are, temperature: 20–25°C, light intensity: 3000–8000 lx, and pH: 7.0–7.5, identified based on orthogonal experiments. The maximum removal rate of total nitrogen, total phosphorus, and chemical oxygen demand was more than 95% in the actual sewage treatment. The sewage treatment capacity of *Desmodesmus* sp. from plateau is higher than that from plains under the same treatment conditions. To identify the differentially expressed genes and metabolites in *Desmodesmus* sp. in response to sewage treatment, a combination of metabolomics and transcriptomics was employed to the microalgae with and without sewage treatment. The results showed that the oxidative phosphorylation, photosynthesis, and propanoate metabolism pathways were the most significantly enriched pathways in sewage treatment. Further, the metabolism of adenosine diphosphate, 2-oxobutanoate, and succinate were significantly upregulated, downregulated, and both upregulated and downregulated, respectively, as shown by the combined transcriptome and metabolome analysis. Additionally, we found that sewage treatment could also induce numerous changes in the primary metabolism, such as carbohydrate, fatty acid biosynthesis, and amino acid metabolism when compared with control. Overall, our results should improve fundamental knowledge of molecular responses to *Desmodesmus* sp. in sewage treatment and contribute to the design of strategies in microalgae response to sewage treatment.

## 1. Introduction

In the recent years, microalgae-based wastewater treatment has attracted increased attention because of its eco-friendly nature and potential economic benefits [1]. Studies have shown that microalgae remove various pollutants from wastewater, such as oxygen consuming pollutants, nitrogen, phosphorus, heavy metals, organic matter, and absorb harmful gases such as NO_X_, SO_X_, and H_2_S at certain concentrations [2]. The biomass harvested from wastewater treatment are also useful as raw materials for biofuel, feed, and chemicals [3-4]. Microalgae can use CO_2_ as the only carbon source for autotrophic growth through photosynthesis and can also use external carbon sources for heterotrophic growth to improve the biomass yield and oil content of microalgae, which is of great significance in reducing global carbon emissions.

Although human society is constantly seeking to improve the quality of spiritual and material life [5], the living environment with industrial development, however, is deteriorating. A large quantity of industrial, aquaculture, and domestic sewage is discharged directly into the natural environment without effective treatment. The sewage is rich in nitrogen, phosphorus, heavy metals, organic matter, etc [6]. Direct discharge of ineffectively treated sewage poses considerable environmental hazards. Currently, the traditional sewage treatment technologies mainly include natural, physical and chemical, biochemical, and the combination of physicochemical and biochemical treatments. However, these traditional treatment processes have disadvantages such as low efficiency, high project and operation costs, and secondary pollution, which result in low penetration rate of various sewage treatment facilities. Therefore, our study aims to find applications of green algae from Yamdrok Lake, Tibet in sewage treatment, which can effectively improve sewage treatment efficiency, reduce cost, energy consumption, and secondary pollution in the existing traditional sewage treatment technology [7].

The principal aims of the present work were: (i) to investigate the best optimal conditions for *Desmodesmus* sp. in wastewater treatment, (ii) to investigate the differences in sewage treatment efficiency of the same *Desmodesmus* sp. obtained from plateau and plains, (iii) to investigate the sewage treatment efficiency of *Desmodesmus* sp. in actual sewage, and (iv) to demonstrate the application of correlation analysis of transcriptomics and metabolomics of *Desmodesmus* sp. used in sewage treatment.

## 2. Materials and Methods

### 2.1 Scale-up cultivation of *Desmodesmus* sp

The culture conditions were as follows: culture duration: 7–15 days, temperature: 25 ± 1°C, light intensity: 3000 lx (photoperiod: 12:12 h), and stirring for 2–3 times every 12 h. Culture volume greater than 1 L was stirred to assist the culture. When microalgae species with long culture period were used, the culture period was appropriately prolonged. After the logarithmic growth phase of *Desmodesmus* sp., i.e., when the final optical density OD_689_ was ≥0.50, the algal culture solution was centrifuged (12000 rpm, 10 min) or filtered (0.45 μm) and was washed thrice with BG-11 culture solution for later use [8].

### 2.2 Determination of the most efficient sewage treatment conditions

One liter algal solution containing *Desmodesmus* sp. was added to 1 L simulated municipal sewage after centrifuging at 12000 rpm for 10 min and were verified by orthogonal experiments with the following parameters. Chemical oxygen demand (COD): 400.0 ± 0.5 mg/L, total nitrogen (TN): 33.0 ± 0.5 mg/L, and total phosphorus (TP): 3.2 ± 0.5 mg/L. Each parameter was tested three times [9].

#### 2.2.1 pH conditions

Experimental conditions to investigate the optimal pH range of wastewater treatment using microalgae are as follows: light intensity: 3000 lx, temperature: 25 ± 1°C, photoperiod: 12:12 h. The initial pH of sewage for treatment was set at 4.0, 7.0, and 10.0, and the sewage treatment was conducted for 7 days while TN, TP, and COD were monitored every 24 h.

#### 2.2.2 Light intensity

Experimental conditions to investigate the optimal range of light intensity for wastewater treatment using microalgae are as follows: temperature: 25 ± 1°C, pH: 7.0 ± 0.5, photoperiod: 12:12 h. The light intensity for sewage treatment was set at 3000 lx, 8000 lx, and 12000 lx, and the sewage treatment was conducted for 7 days while TN, TP, and COD were monitored every 24 h.

#### 2.2.3 Temperature conditions

Experimental conditions to investigate the optimal temperature range of microalgae wastewater treatment are as follows: light intensity: 3000 lx, pH: 7.0 ± 0.5°C, and photoperiod: 12:12h. The temperature gradient for sewage treatment was set at 10, 20, 30, and 40°C, and the sewage was treated continuously for 7 days while the TN, TP, and COD were monitored every 24 h.

### 2.3 Application of *Desmodesmus* sp. in actual sewage treatment

The optimal sewage treatment conditions for microalgae were determined from orthogonal experiments and were employed for the actual sewage treatment. Continuous sewage treatment was conducted for 7 days. TN, TP, and COD were monitored every 24 h. Domestic sewage from the sewage outlet of a college was collected as the experimental sample, and *Desmodesmus* sp. was used to assess the sewage treatment effect. The actual sewage parameters are as follows: COD 150.0 mg/L, TN 42.5 mg/L, and TP 1.5 mg/L. Each parameter was tested three times.

### 2.4 Sewage treatment using *Desmodesmus* sp. obtained from plateau and plains

Positive control samples were purchased from Wuhan Institute of Aquatic Sciences, sampled from Taihu Lake, Jiangsu Province (product number: FACHB-2921), which were reserved by amplification culture.

*Desmodesmus* sp. was isolated from Yamdrok Lake, Shannan City, Tibet Autonomous Region. The sample was separated, purified, and cultured by amplification in the laboratory for future use.

One liter of algal solution with the same absorbance was selected and filtered through 0.45 µm or centrifuged at 12000 rpm for 10 min to obtain the filtered or centrifuged algae, respectively, which was then introduced to the simulated municipal sewage for sewage treatment. The experimental conditions are as follows: temperature gradient: 25 ± 1°C, light intensity: 5000 lx, pH: 7.0 ± 0.5, photoperiod: 12:12 h, and continuous sewage treatment for 7 days. TN, TP, and COD were monitored every 24 h.

### 2.5 RNA extraction, cDNA library construction, and RNA-sequencing

#### 2.5.1 RNA quantification and integrity assessment

By using RNA Nano 6000 Assay Kit of the Bioanalyzer 2100 system (Agilent Technologies, CA, USA), total amount and integrity of RNA were assessed. For all RNA-Seq experiments from microalgae before and after sewage treatment, three biological replicates were used [10].

#### 2.5.2 Library preparation for transcriptome sequencing

poly-T oligo-attached magnetic beads were used to purify mRNA from total RNA. The library fragments were purified using the AMPure XP system (Beckman Coulter, Beverly, USA) to obtain the best cDNA fragments with a length of 370–420 bp [11-12]. The PCR product was purified using AMPure XP beads to obtain the library. In order to ensure the quality of the library, firstly, the library was analyzed quantitatively by Qubit 2.0 Fluorometer; Then, the insertion size of the library was detected by using Agilent 2100 biological analyzer; Finally, the effective concentration of the library was accurately quantified by using QRT PCR (the effective concentration of the library was higher than 2 nm) [13].

#### 2.5.3 Clustering and sequencing

The basic principle of sequencing is synthetic sequencing, that is, synthesis and sequencing at the same time. Amplification was performed by adding four prepared fluorescent labeled dNTP, DNA polymerase, and splicing primers to the sequenced flow cytometry cells. When the sequence cluster extended the complementary chain, the fluorescently labeled dNTP was sequenced using Illumina novaseq 6000, resulting in a final reading of 150 bp pairs [14].

The raw sequence reads were deposited in the NCBI Sequence Read Archive (http://www.ncbi.nlm.nih.gov/subs/sra, accession number PRJNA830552) (accessed on 22 April 2022).

### 2.6 De novo assembly, annotation, and classification

#### 2.6.1 Quality control

Firstly, the image data measured by the high-throughput sequencer were converted into sequence data (reads) by CASAVA base recognition. Then, the raw data (raw reads) in fastq format were processed by the internal Perl script to obtain clean data (clean read), and the Q20, q30 and GC contents of clean data are calculated at the same time. The obtained high-quality cleaning data are used for downstream analysis [15-16].

#### 2.6.2 Transcriptome assembly and quality evaluation

The cleaning data obtained were analyzed using the Trinity software package (v2.6.6), and then the cleaning readings of the reference sequence components were obtained. Transcriptome assembly is completed on the left [17]. Busco software was used to evaluate the splicing quality of Trinity and the accuracy and integrity of splicing results [17].

#### 2.6.3 Corset hierarchical clustering and Gene functional annotation

Corset (version 4.6) was used to aggregate transcripts into many new clusters, which were defined as “Gene” [18]. Gene functions were annotated through the following databases: Nr (NCBI non-redundant protein sequences), Nt (NCBI non-redundant nucleotide sequences), Pfam (Protein family), KOG/COG (Clusters of Orthologous Groups of proteins), Swiss-Prot (A manually annotated and reviewed protein sequence database), KO (KEGG Ortholog database), and GO (Gene Ontology) [19-20].

### 2.7 Differential gene expression analysis and enrichment analysis

#### 2.7.1 Differential expression analysis

Differential expression analysis was performed using DESeq2 R package (v1.20.0). The resulting P-values were adjusted using the Benjamini–Hochberg method to control the error detection rate. Padj < 0.05 and |log2(foldchange)| > 1 were set as thresholds for significant differential expression [21].

#### 2.7.2 GO and KEGG enrichment analysis of differentially expressed genes

Go function and KEGG pathway enrichment analysis of differential gene sets are realized by GOseq (v1.10.0) and KOBAS (v2.0.12) software. All differential gene sets, upregulated differential gene sets and downregulated differential genes enriched in each differential comparison combination were reflected in the results of enrichment analysis [22].

### 2.8 Untargeted metabolome analysis

#### 2.8.1 Metabolite extraction

The analysis was performed using a liquid chromatography tandem mass spectrometry (LC-MS/MS) system. Sample preparation: firstly, the samples were resuspended with precooled 80% methanol through eddy current, then melted and rotated on ice for 30 s and treated with sonication for 6 min, and finally centrifuged at 5000 rpm and 4 °C for 1 min. The supernatant was freeze-dried and dissolved with 10% methanol for standby [23].

#### 2.8.2 Metabolite analysis

Ultra-high performance (UHP) LC-MS/MS analyses were performed using a Vanquish UHPLC system (ThermoFisher, Germany) coupled with an Orbitrap Q Exactive™ HF mass spectrometer (Thermo Fisher, Germany) at Novogene Co., Ltd. (Beijing, China). Raw data files generated by UHPLC-MS/MS were processed using a compound discoverer 3.1 (CD3.1, ThermoFisher) for peak alignment, peak pickup and quantification of each metabolite. Statistical software R (VR-3.4.3), Python (V2.7.6) and CentOS (CentOS 6.6) were used for statistical analysis. For non-normally distributed data, the area normalization method is used to attempt normal transformation [24]. Try to perform normal transformation on the data with non normal distribution through the area normalization method [24].

### 2.9 Metabolomics data analysis

The metabolites were annotated using KEGG (https://www.genome.jp/kegg/pathway.html), HMDB (https://hmdb.ca/metabolites), and LIPIDMaps databases (http://www.lipidmaps.org/). The analysis of principal component analysis (PCA) and partial least squares discriminant analysis (PLS-DA) were realized by metaX software. Statistical significance (P-value) was calculated by univariate analysis (t-test). The criteria of differential metabolites were VIP > 1, P-value < 0.05, folding metabolites ≥ 2 or ≤ 0.5. Metabolites of interest were used to filter by volcano plots, which was based on log 2 (fold change) and -log 10 (P-value) of metabolites by ggplot2 in R language [25]. The data were normalized by using the Z-scores of the intensity region of differential metabolites, and the clustering heat maps were plotted by using the Phatmap software package in R language. In R, the Pearson correlation coefficient between different metabolites were analyzed by cor (). When P-value < 0.05, it was considered to be statistically significant. The functions and metabolic pathways of metabolites were compared by using KEGG database. When the ratio of X / N > y / N were satisfied, the corresponding metabolic pathways were considered to be enriched. When P-value < 0.05, metabolic pathway enrichment was considered statistically significant.

### 2.10 Combined transcriptome and metabolome analyses

The transcription-metabolite network was constructed from gene metabolite networks with Pearson correlation coefficient > 0.8. All differential genes and metabolites obtained were mapped to the KEGG pathway database at the same time, so as to determine the main biochemical and signal transduction pathways involved in differential metabolites and genes [26].

## 3. Results

### 3.1 Optimal pH conditions for wastewater treatment

At pH 7.0, the most efficient treatment of COD was 97.8% on the third day. Similarly, at pH 4.0, the most efficient COD treatment was 96.3% on the fourth day. Further, at pH 10.0, the most efficient COD treatment was 97.0% on the third day (Fig. 1a). Considering the stability of sewage treatment index, the effective COD treatment was in the following order: pH 7.0 > pH 10.0 > pH 4.0.

**Fig. 1.**
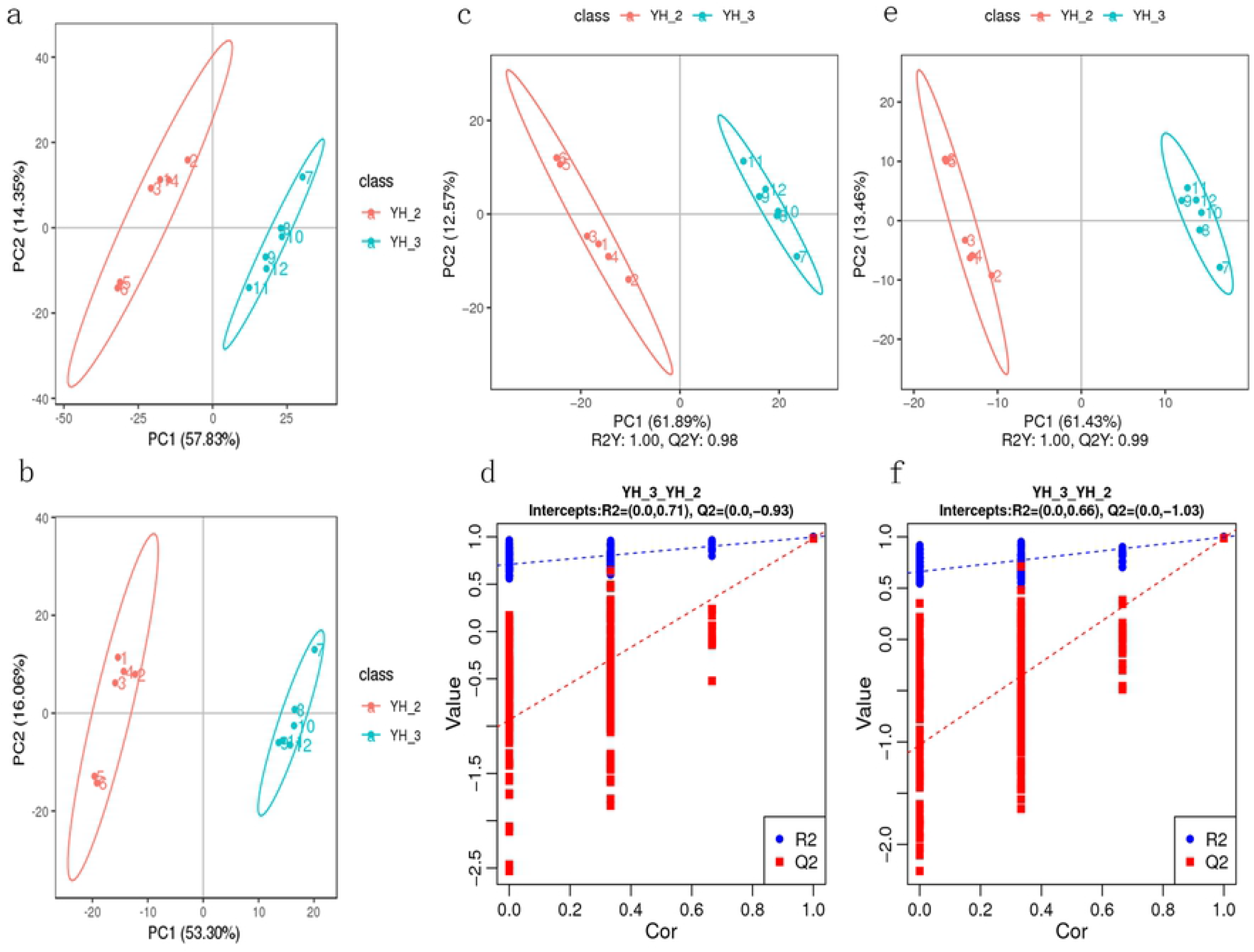
Orthogonal test verification of simulated municipal sewage treatment by Tibetan *Desmodesmus* sp.: (a) COD treatment trends under different pH conditions; (b) TP treatment trends under different pH Conditions; (c) TN treatment trends under different pH conditions; (d) COD treatment trends under different light intensities; (e) TP treatment trends under different light intensities; (f) TN treatment trends under different light intensities; (g) COD treatment trends under different temperatures; (h) TP treatment trends under different temperatures; (i) TN treatment trends under different temperatures.

At neutral pH 7.0, the most efficient TP treatment was 99.4% on the third day. At the acidic pH 4.0, the most efficient TP treatment was 49.1% on the third day. At the alkaline pH 10.0, the most efficient TP treatment was 65.6% on the third day (Fig. 1b). Considering the stability of sewage treatment index, the effective TP treatment was in the following order: pH 7.0 > pH 10.0 > pH 4.0.

At neutral pH 7.0, the most efficient TN treatment was 99.6% on the third day. At the acidic pH 4.0, the most efficient TN treatment was 94.3% on the second day. At the alkaline pH 10.0, the most efficient TN treatment was 96.2% on the third day (Fig. 1c). Considering the stability of sewage treatment index, the effective TN treatment was in the following order: pH 7.0 > pH 10.0 > pH 4.0.

### 3.2 Optimal light intensity for wastewater treatment

At 3000 lx light intensity, the most efficient COD treatment was 96.1% on the second day. At 8000 lx light intensity, the most efficient COD treatment was 98.2% on the second day. At 12000 lx light intensity, the most efficient COD treatment was 95.4% on the third day (Fig. 1d). The efficiency of COD treatment was in the following order: 8000 lx > 3000 lx > 12000 lx.

At 3000 lx light intensity, the most efficient TP treatment was 90.9% on the fifth day. At 8000 lx light intensity, the most efficient TP treatment was 89.4% on the second day. At 12000 lx light intensity, the most efficient TP treatment was 72.8% on the third day (Fig. 1e). Although the best efficiency was 90.9% under 3000 lx light intensity, the 89.4% treatment efficiency at 8000 lx was of the shortest duration. Thus, the efficiency of TP treatment was of the order 8000 lx > 3000 lx > 12 000 lx when the sewage treatment duration is considered.

At 3000 lx, the most efficient TN treatment was 97.0% on the fifth day. At 8000 lx light intensity, the most efficient TN treatment was 99.3% on the second day (Fig. 1f). At 12000 lx, the most efficient TN treatment was 96.7% on the first day. The effective light intensity for TN treatment was in the following order: 8000 lx > 3000 lx > 12000 lx.

### 3.3 Optimal temperature conditions for wastewater treatment

At 10°C, the most efficient COD treatment was 93.5% on the second day. At 20°C, the most efficient COD treatment was 96.1% on the second day. At 30°C, the most efficient COD treatment was 96.0% on the second day. At 40°C, the most efficient COD treatment was 94.6% on the second day (Fig. 1g). The effective temperature for COD treatment was in the following order: 20°C > 30°C > 40°C > 10°C.

At 10°C, the most efficient TP treatment was 83.8% on the fourth day. At 20°C, the most efficient TP treatment was 89.7% on the fifth day. At 30°C, the most efficient TP treatment was 81.3% on the fifth day. At 40°C, the most efficient TP treatment was 87.5% on the fifth day (Fig. 1h). The effective temperature for TP treatment was in the following order 20°C > 30°C > 40°C > 10°C.

At 10°C, the most efficient TN treatment was 76.9% on the fourth day. At 20°C, the most efficient TN treatment was 95.0% on the fourth day. At 30°C, the most efficient TN treatment was 86.4% on the fourth day. At 40°C, the most efficient TN treatment was 82.7% on the third day (Fig. 1i). The effective temperature for TN treatment was of the order 20°C > 30°C > 40°C > 10°C.

### 3.4 Application of Desmodesmus sp. in actual sewage treatment

Based on the above orthogonal experiments, the optimal sewage treatment conditions for *Desmodesmus* sp. from Yamdrok Lake are as follows: temperature: 20–25°C, light intensity: 3000–8000 lx, and pH: 7.0. Under these conditions, the experimental conditions are as follows: temperature: 25 ± 1°C, light intensity: 4000 lx, and pH: 7.0 ± 0.5.

From the treatment of actual sewage using *Desmodesmus* sp., the most efficient COD treatment was 93.4% on the third day (Fig. 2a). Similarly, the most efficient TP treatment was 99.3% on the second day (Fig.2b). Further, the most efficient TN treatment was 98.7% on the third day (Fig. 2c).

**Fig. 2.**
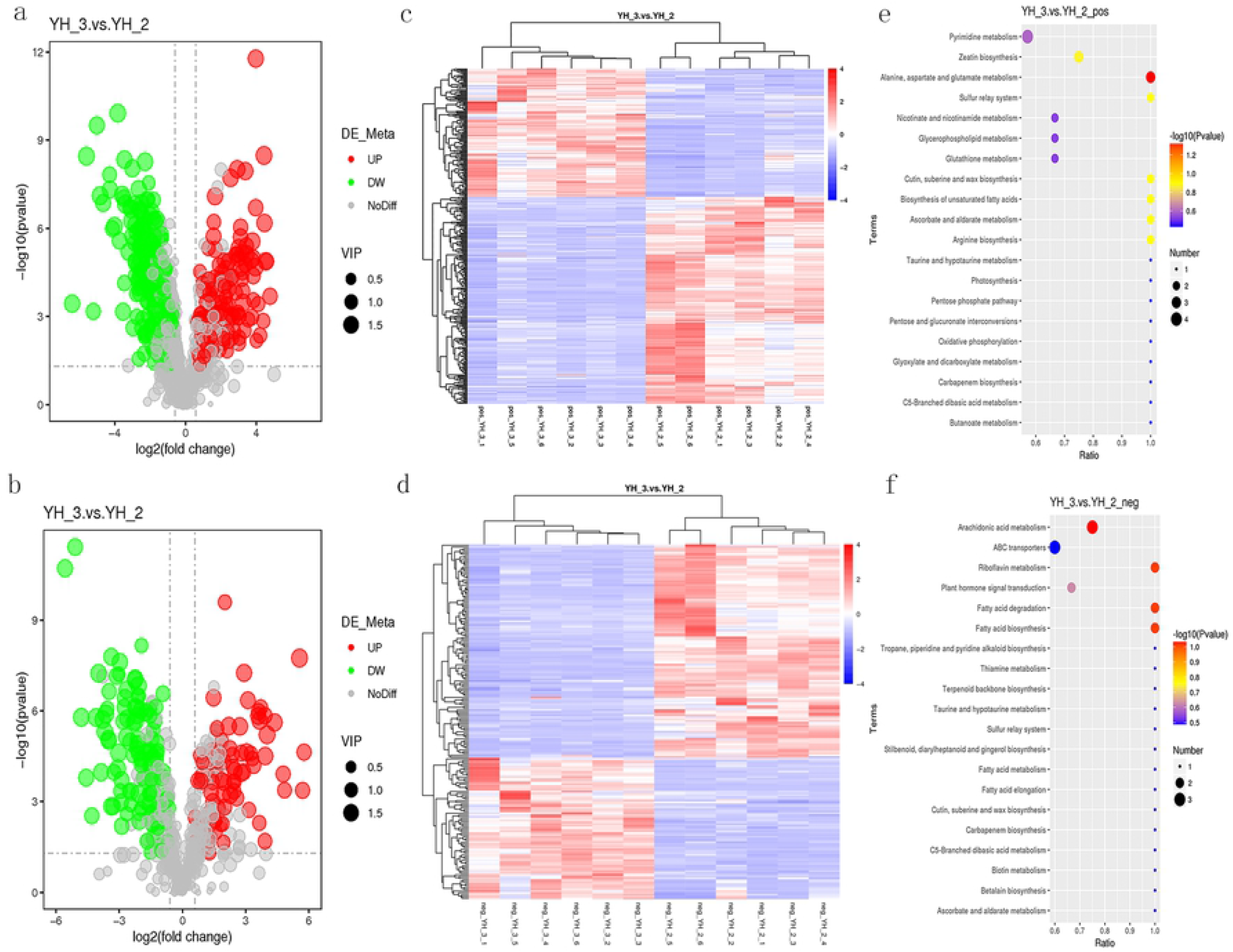
Application of *Desmodesmus* sp. in actual sewage treatment, and comparative experiment of sewage treatment. (a) COD treatment trends; (b) TP treatment trends; (c) TN treatment trends. Positive control experiment of *Desmodesmus* sp. from plateau and plain areas: (d) COD treatment trends; (e) TP treatment trends; (f) TN treatment trends.

### 3.5 Positive control experiment of Desmodesmus sp. collected from different altitudes

Based on the optimal sewage treatment conditions, *Desmodesmus* sp. from plateau area reduced COD on the fourth day of treatment, and the efficiency was 97.0%. *Desmodesmus* sp. from plains reduced COD on the fourth day of treatment, and the efficiency was 91.9% (Fig. 2d). The effective COD treatment is of the order *Desmodesmus* sp. from the plateau > *Desmodesmus* sp. from the plains.

*Desmodesmus* sp. from plateau reduced the TP on the second day of treatment by 95.5% and those from plains reduced the TP on the fifth day of treatment by 89.7% (Fig. 2e). The effective TP treatment is of the order *Desmodesmus* sp. from the plateau > *Desmodesmus* sp. from the plains.

*Desmodesmus* sp. from plateau reduced the TN on the third day of treatment by 99.1%, whereas *Desmodesmus sp*. from plains reduced TN on the third day of treatment by 86.1% (Fig. 2f). The effective TN treatment was of the order *Desmodesmus sp*. from the plateau > *Desmodesmus sp*. from the plains.

Under the same treatment conditions, the sewage treatment capacity of *Desmodesmus sp*. from plateau area was higher than that from plains area.

### 3.6 RNA sequencing and differential gene expression analysis in non-parametric transcriptomics

Transcription and widely targeted metabolite profiles of *Desmodesmus* sp. that were employed for (YH_3) and not employed (YH_2) for sewage treatment were explored. Three independent biological replicates were used for each treatment, resulting in six samples. A transcriptome database containing 217 942 unigenes of average length 1106 bp was obtained using Trinity software, with an N50 and N90 lengths of 1578 bp and 475 bp, respectively.

All unigenes and transcripts obtained by transcriptome assembly were aligned with five major databases (NR, NT, KOG, PFAM, and GO). A total of 13633 assembled unigenes were found to have homologs in the databases NR (119989), NT (26206), KOG (49371), PFAM (123051), and GO (123041) (Fig. 3a). The transcriptome changes of *Desmodesmus* sp. were investigated through RNA-Seq analysis. More than 20 million reads were generated per sample. Of these reads, the Q30 percentage (sequencing error rate < 1%) was over 92%, and GC content was approximately 58% for the libraries. Among all the libraries, 77.72–82.87% of unique reads were mapped to the *Desmodesmus* sp. genome (Table 1). A total of 18872 and 35871 genes in sewage treatment were significantly upregulated and downregulated, respectively (DESeq2 padj < 0.05 |log2FoldChange| > 1, Fig. 3b).

**Table 1.**
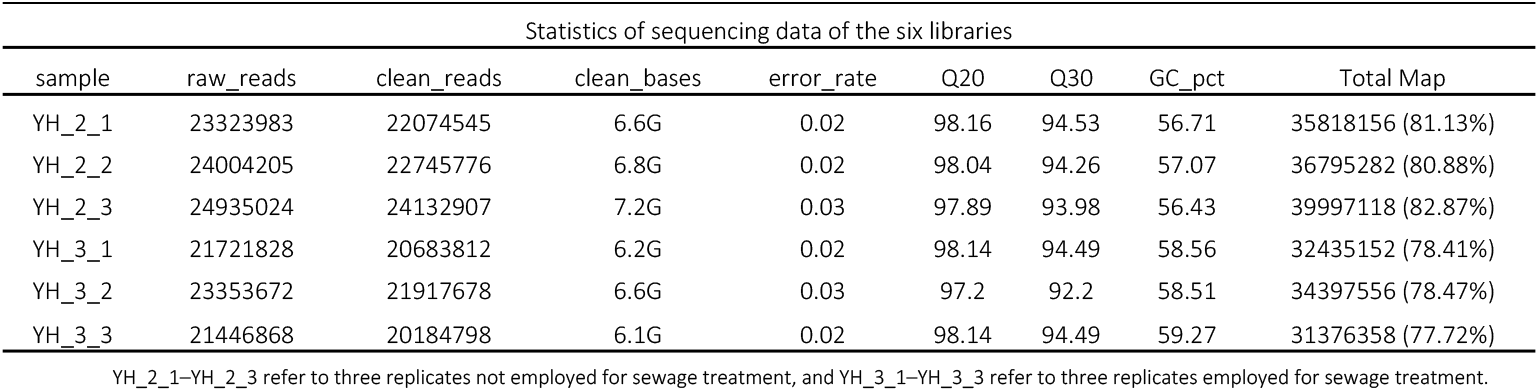
Statistics of sequencing data of all libraries.

| Statistics of sequencing data of the six libraries |  |  |  |  |  |  |  |  |
| --- | --- | --- | --- | --- | --- | --- | --- | --- |
| sample | raw_reads | clean_reads | clean_bases | error_rate | Q20 | Q30 | GC_pct | Total Map |
| YH_2_1 | 23323983 | 22074545 | 6.6G | 0.02 | 98.16 | 94.53 | 56.71 | 35818156 (81.13%) |
| YH_2_2 | 24004205 | 22745776 | 6.8G | 0.02 | 98.04 | 94.26 | 57.07 | 36795282 (80.88%) |
| YH_2_3 | 24935024 | 24132907 | 7.2G | 0.03 | 97.89 | 93.98 | 56.43 | 39997118 (82.87%) |
| YH_3_1 | 21721828 | 20683812 | 6.2G | 0.02 | 98.14 | 94.49 | 58.56 | 32435152 (78.41%) |
| YH_3_2 | 23353672 | 21917678 | 6.6G | 0.03 | 97.2 | 92.2 | 58.51 | 34397556 (78.47%) |
| YH_3_3 | 21446868 | 20184798 | 6.1G | 0.02 | 98.14 | 94.49 | 59.27 | 31376358 (77.72%) |

YH\_2\_1–YH\_2\_3 refer to three replicates not employed for sewage treatment, and YH\_3\_1–YH\_3\_3 refer to three replicates employed for sewage treatment.

**Fig. 3.**
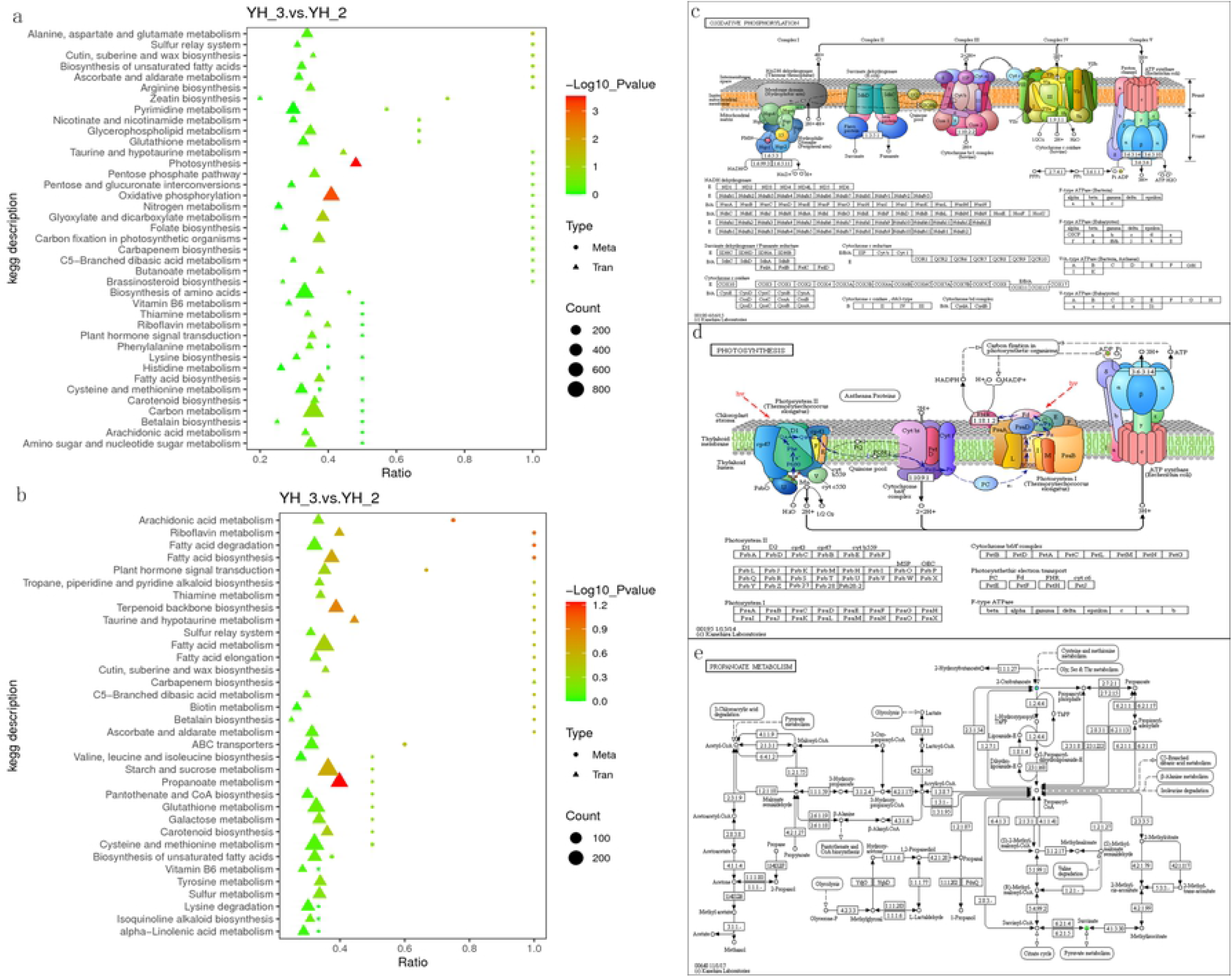
(a) Functional annotation numbers of unigenes in the NR, NT, KOG, PFAM and GO databases; (b) Number of DEGs; (c) GO classification; (d) KEGG enrichment of DEGs.

All the DEGs were subjected to GO and KEGG analysis. On comparing YH_2 with YH_3, we found that the significantly enriched GO terms (p < 0.01) were linked to biosynthetic, small molecule metabolic, and cellular protein modification processes in the “biological process” category. Cytoplasm was the most enriched in the “cellular component” category. Ion binding and oxidoreductase activity were the most enriched in the “molecular function” category (Fig. 3c, Fig. S1). Pathways showing significant change (p < 0.01) in sewage treatment were identified using the KEGG database, the DEGs were involved in three enriched pathways: ribosome, oxidative phosphorylation, and ubiquitin-mediated proteolysis (Fig. 3d, Fig. S2).

### 3.7 Differential metabolite accumulation analysis related to Desmodesmus sp. employed for sewage treatment

PCA of the DEGs and differentially accumulated metabolites showed that the YH_3 treatment group showed obvious differences with the YH_2 (control) group, which explained 57.83% and 53.30% of the total variation (Fig. 4a, b). These results indicated that *Desmodesmus* sp. has good treatment effect on sewage.

**Fig. 4.**
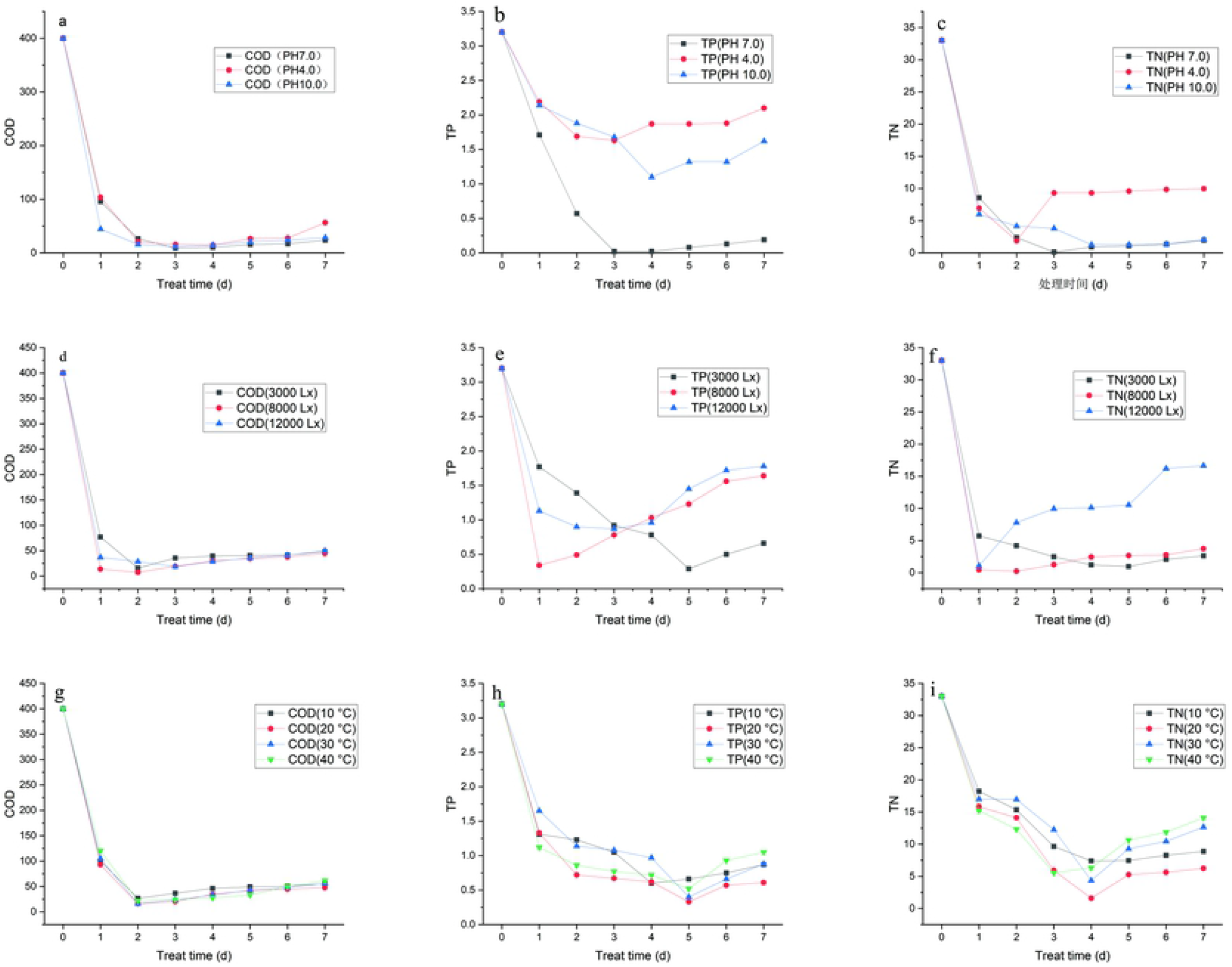
PCA in YH_3 vs. YH_2. (a) PCA of the variance-stabilized estimated raw counts of differentially accumulated metabolites in positive polarity mode. (b) PCA of the variance-stabilized estimated raw counts of differentially accumulated metabolites in negative polarity mode. (c) PLS-DA of differentially accumulated metabolites in positive polarity mode. (d) Permutation test for positive polarity mode. (e) PLS-DA of differentially accumulated metabolites in negative polarity mode. (f) Permutation test for negative polarity mode.

In the scattered plots, abscissa is the score of the sample on the first principal component; ordinate is the score of the sample on the second principal component. R2Y represents the interpretation rate of the model, and Q2Y is used to evaluate the prediction ability of the Partial Least Squares Discrimination (PLS-DA) model. The PLS-DA models of each comparison group were established, and the model evaluation parameters (R2, Q2) were obtained by seven-fold cross validation (when the biological repetition number of the sample n ≤ 3, it was K cycles of interactive verification, k = 2n). The closer R2 and Q2 are to 1, the more stable and reliable is the model, when R2Y is greater than Q2Y, the model is well established (Fig. 4c, e). The grouping marks of each sample are randomly disrupted before modeling and prediction. Each modeling corresponds to a set of R2 and Q2 values. Their regression lines can be obtained according to the Q2 and R2 values after 200 disruptions and modeling. When R2 data is greater than Q2 data and the intercept between Q2 regression line and Y-axis is less than 0, it indicates that the model is not “over-fitting,” suggesting that the model can better describe the sample and can be used as the premise for the search of model biomarker group (Fig. 4d, f).

Untargeted metabolomics was applied for the evaluation of differentially accumulated metabolites in YH_3 and YH_2. After sewage treatment under the positive polarity mode, 910 metabolites were identified, of which 366 were significantly different, 140 were significantly upregulated, and 226 were significantly downregulated (YH_3 vs. YH_2, VIP > 1.0, FC > 1.5 or FC < 0.667, and padj < 0.05, Fig. 5a, Table S1). Under negative polarity mode, 543 metabolites were identified, 196 of which were significantly different, 78 were significantly upregulated, and 118 were significantly downregulated (YH_3 vs. YH_2, VIP > 1.0, FC > 1.5 or FC < 0.667, and padj < 0.05, Fig. 5b, Table S2).

**Fig. 5.**
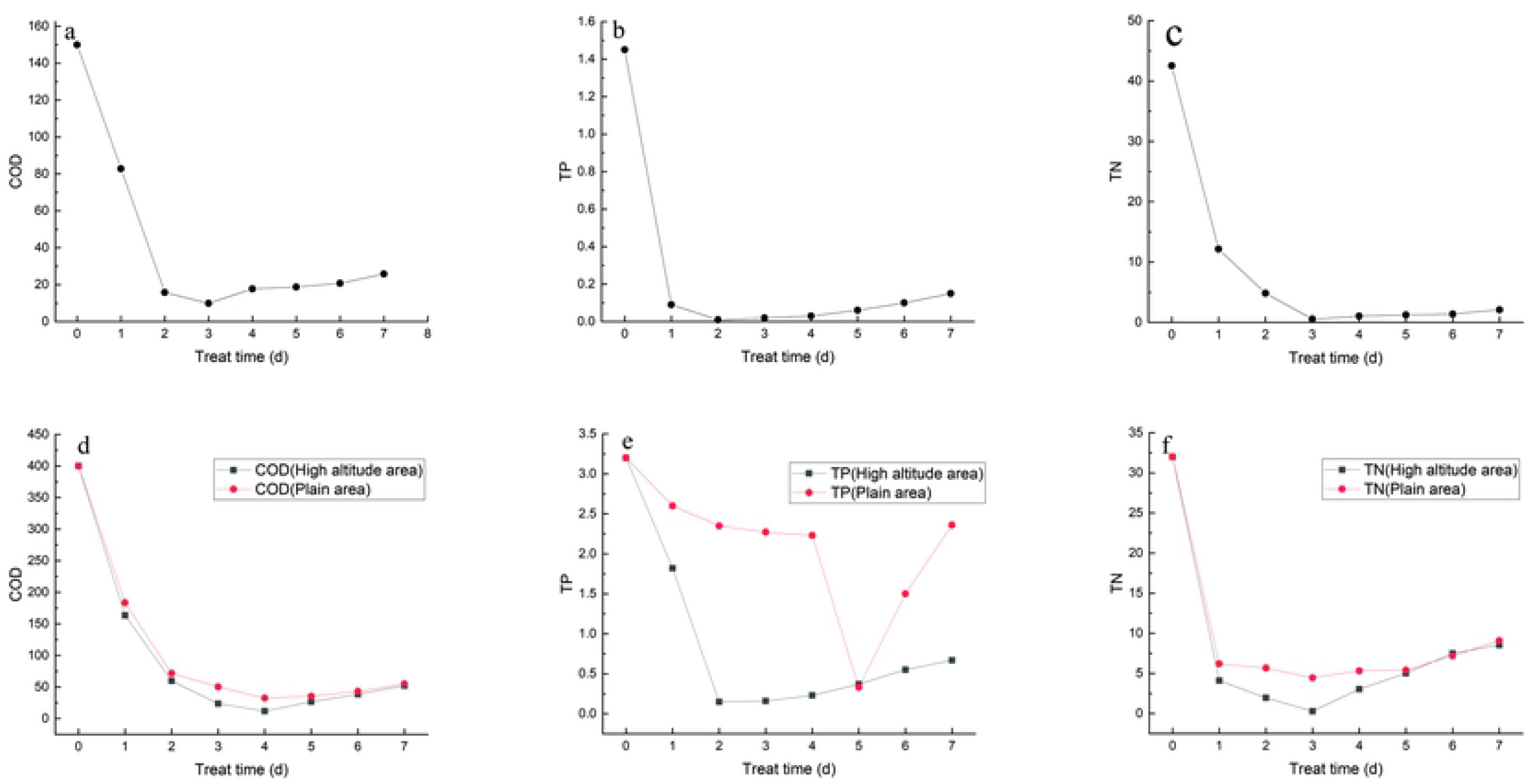
Differential metabolite accumulation analysis. (a) Volcano plot of differential metabolites in positive polarity mode. (b) Volcano plot of differential metabolites in negative polarity mode. Each point in the volcano map represents a metabolite, X-axis represents the logarithm of the quantitative difference of a certain metabolite in the two samples, and Y-axis represents the difference significance level (-log10p value). The green dots represent downregulated differentially accumulated metabolites, the red dots represent upregulated differentially accumulated metabolites, and the gray represents detected but not significantly differentially accumulated metabolites. YH_2: *Desmodesmus* sp. without sewage treatment; YH _3: *Desmodesmus* sp. with sewage treatment. (c) HCA of differentially accumulated metabolites in YH_3 vs. YH_2 (6 replicates for each sample) by positive polarity mode. (d) HCA of differentially accumulated metabolites in YH_3 vs. YH_2 (6 replicates for each sample) by negative polarity mode. (e) Scatter plot of KEGG pathways enriched in positive polarity mode. (f) Scatter plot of KEGG pathways enriched in negative polarity mode. Abscissa in the figure is the ratio of the number of differential metabolites in the corresponding metabolic pathway to the number of total metabolites identified in the pathway. The greater the value, the higher is enrichment degree of differential metabolites in the pathway. The color of the point represents the p-value value of the hypergeometric test. The smaller the value, the greater is the reliability and statistical significance. The size of the point represents the number of differential metabolites in the corresponding pathway. The larger the point, the more is the number of differential metabolites in the pathway.

For the hierarchical cluster analysis (HCA) of differentially accumulated metabolites in YH_3 and YH_2 (6 replicates for each sample), we found that there were significant differences in metabolites of *Desmodesmus* sp. before and after sewage treatment (Fig. 5c, d). Furthermore, for differentially accumulated metabolites in positive polarity mode, we found that the differentially accumulated metabolites were most significantly enriched in KEGG pathways to alanine, aspartate, and glutamate metabolism (Fig. 5e).

For differentially accumulated metabolites in negative polarity mode, we found that the differentially accumulated metabolites were most significantly enriched in KEGG pathways to arachidonic acid metabolism, fatty acid biosynthesis, fatty acid degradation, and riboflavin metabolism (Fig. 5f).

### 3.8 Combined transcriptome and metabolome analyses

To quantitatively map the transcripts directly to metabolic pathways involved in *Desmodesmus* sp. after sewage treatment, the co-joint KEGG pathway enrichment analysis of transcriptome and metabolome was performed. The results showed that the same pathways of DEGs and differentially accumulated metabolites were enriched to oxidative phosphorylation and photosynthesis (p < 0.01) in positive polarity mode (Fig. 6a), and propanoate metabolism (p < 0.01) in negative polarity mode (Fig. 6b). To better understand the relationship between genes and metabolites, the DEGs and differentially accumulated metabolites were simultaneously mapped to the KEGG pathway diagram (Table S3, Table S4). As shown in Fig. 6c and 6d, the metabolites of adenosine diphosphate were simultaneously mapped to the oxidative phosphorylation (ko00190) and photosynthesis (ko00195) in positive polarity mode. Further, two differentially accumulated metabolites (2-oxobutanoate and succinate) were simultaneously mapped to the propanoate metabolism (ko00640) in negative polarity mode.

**Fig. 6.**
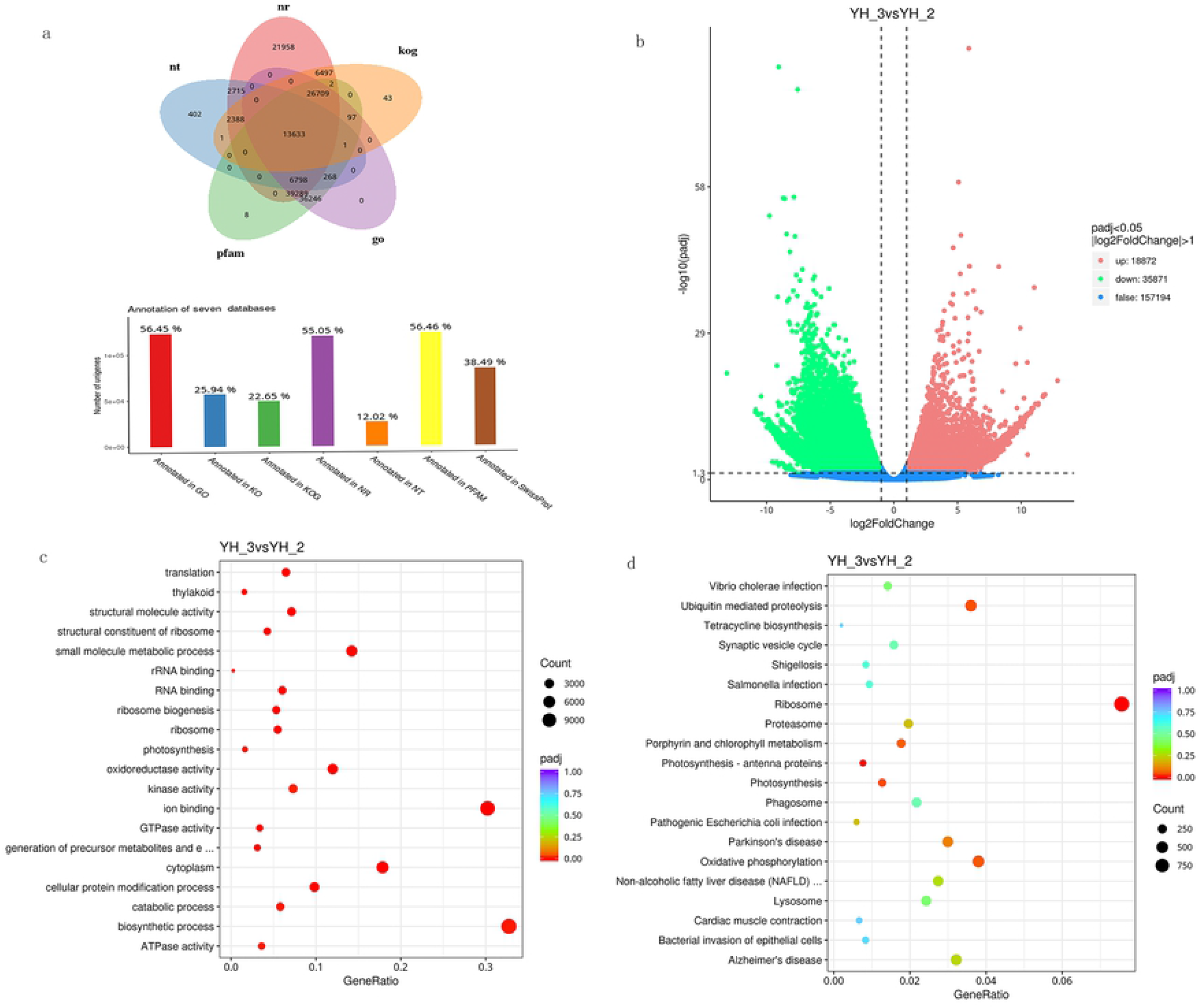
Joint analysis of the differentially expressed genes and differentially accumulated metabolites in YH_3 vs. YH_2. (a) Scatter plot of differential genes and metabolites in positive polarity mode. (b) Scatter plot of differential genes and metabolites in negative polarity mode. (c) Differentially expressed genes and differentially accumulated metabolites simultaneously mapped to the oxidative phosphorylation (ko00190). (d) Differentially expressed genes and differentially accumulated metabolites simultaneously mapped to the photosynthesis (ko00195). (e) Differentially expressed genes and differentially accumulated metabolites simultaneously mapped to the propanoate metabolism (ko00640). Solid green circles indicate the metabolites noted, red circles indicate a significant gene or metabolite upregulation, blue circles indicate a significant gene or metabolite downregulation of the, and solid circles in green represents a gene that is both upregulated and downregulated.

## 4. Discussion

Microalgae are valuable microorganisms and play an increasingly important role in industrial, agricultural, and domestic wastewater treatment. However, there are few reports on the application of microalgae in sewage treatment in the Qinghai–Tibet Plateau. Thus, it was deemed necessary to find a suitable microalga to treat municipal sewage on the plateau [27]. Various high-quality omics techniques have been reported, which may deepen our understanding of the response of microalgae to sewage treatment. In this study, we studied the DEGs and differentially accumulated metabolites based on combined transcriptomic and metabolomic profiling of YH_3 and YH_2 of *Desmodesmus* sp. to explore the sewage treatment mechanisms.

### 4.1 Optimum conditions of Tibetan *Desmodesmus sp*. for sewage treatment

The Qinghai-Tibet Plateau is marked by a unique natural environment with high altitude, low oxygen, low pressure, and strong ultraviolet radiation [28]. Microalgae have adapted to the extreme environment of the plateau and are pivotal in maintaining the water quality of the plateau. Therefore, we aimed at finding the best sewage treatment conditions for microalgae from high-altitude areas and establishing a reliable technical condition for subsequent practical application.

We optimized and determined the optimal treatment conditions of *Desmodesmus* sp. for sewage treatment, and the specific treatment conditions include: treatment temperature: 20–25°C, light intensity: 3000–8000 lx, and the pH of the reaction system: 7.0–7.5. The most efficient treatment of COD, TP, and TN was on the third day, and the efficiency was more than 95.0%. Particularly, we found that Tibetan *Desmodesmus* sp. are efficient in treating sewage under light intensity of approximately 8000 lx, at a low temperature of 10°C, under acidic and alkaline conditions, indicating that they are adaptable to extreme environment in high altitude areas. Thus, they show good tolerance to sewage that supports microbial growth. In the later stage, the metabolism and cell lysis of microalgae continued. Phosphate and organic matter were released in these processes, and the organic compounds released into the treated sewage were hydrolyzed into easily biodegradable organic matter, which finally led to the rise of corresponding indicators because of nutrient shortage in simulated sewage [29]. Therefore, in the actual sewage treatment process it is necessary to timely control the end point of sewage treatment by harvesting the microalgae in time [30].

### 4.2 Positive control test

To further verify the efficiency of Tibetan *Desmodesmus* sp. in actual domestic sewage treatment and the sewage treatment efficiency of the same genus as that in plain areas, we performed a positive control test. In the experiment, the domestic sewage discharged from a college was used for complete sewage treatment, and the effective removal rates of TN, TP, and COD was more than 95%. Under the parallel experimental conditions, the efficiency of Tibetan *Desmodesmus* sp. to treat sewage was higher than that of the same genus from Taihu Lake. This is because of the adaptation of *Desmodesmus* sp. in high-altitude areas to low oxygen, low pressure, low temperature, and high radiation conditions. Thus, it has stronger adaptability in the sewage treatment processes.

### 4.3 Changes in primary metabolites related to sewage treatment

Microalgae are known to produce economically beneficial substances such as amino acids, polysaccharides, carbohydrates, proteins, and fatty acids during sewage treatment processes [31]. In this study, we found that the differentially accumulated metabolites were most significantly enriched in KEGG pathways to alanine, aspartate, glutamate, arachidonic acid, and riboflavin metabolisms, fatty acid biosynthesis, and fatty acid degradation. Furthermore, we found the key genes related to these processes (Table S5). Notably, the upregulated metabolites were octadecenoic acid and adenylosuccinate, and the both upregulated and downregulated metabolites were PGA2, 16(R)-HETE, hexadecenoic acid, riboflavin, L-asparagine, and L-glutamate, which might play a crucial role in carbohydrate, fatty acid, and amino acid biosynthesis in sewage treatment. These results reveal that primary metabolite (carbohydrate metabolism, fatty acid biosynthesis metabolism, and amino acid metabolism) pathways play important roles in sewage treatment [32].

### 4.4. Signaling pathways related to sewage treatment

In this study, the transcriptional and metabolic association analysis revealed that metabolism of adenosine diphosphate was significantly upregulated by key genes related to photosynthesis and oxidative phosphorylation (Table S5). Our results showed that sewage treatment activated these genes, which are the key genes in adenosine-triphosphate (ATP) biosynthesis [33]. ATP plays an important role in nucleic acid synthesis, and it stores and transmits chemical energy and provides energy for life activities. The direct source of energy for life activities is ATP, which participates in microalgae photosynthesis, nutrient transport (active transport consumes energy), and cell division and provides energy to microalgae for sewage treatment. Our results indicated that the photosynthesis and oxidative phosphorylation signaling pathways were the most enriched pathways which play important roles in *Desmodesmus* sp. response to sewage treatment. We found that many genes involved in energy metabolism were induced by *Desmodesmus* sp. response to sewage treatment including oxidative phosphorylation and carbon fixation in photosynthetic organisms and carbohydrate metabolism [34].

### 4.5 Secondary metabolite pathway related to *Desmodesmus* sp. response to sewage treatment

In our study, the results of both transcriptome and metabolome analyses showed that propanoate metabolism pathways were induced by sewage treatment and key genes related to propanoate metabolism (Table S5). Furthermore, the downregulated metabolite was 2-oxobutanoate (2-OBA), which as a toxic metabolic intermediate, generally arrests the cell growth of most microorganisms and blocks the biosynthesis of target metabolites [35-36]. In the wastewater treatment process, the metabolite of 2-OBA is downregulated, which can promote the effect of microalgae on wastewater treatment and maintain the normal metabolism of microalgae. The upregulated and downregulated metabolite was succinate, which might play an important role in regulating pyruvate metabolism. In the metabolic process, pyruvate is the final product of glycolysis pathway. It enters mitochondria for oxidation to produce acetyl CoA, enters the tricarboxylic acid cycle and is oxidized to carbon dioxide and water to complete the aerobic oxidation and energy supply process of glucose. Thus, maintaining cell viability of microalgae plays an important role in the absence of nutrients or light. Pyruvate can also realize the mutual transformation among sugar, fat, and amino acids through acetyl CoA and tricarboxylic acid cycle. Therefore, pyruvate plays an important role in the metabolic connection of the three nutrients and maintains the normal metabolism of microalgae [37]. Therefore, the induction of propanoate metabolism related genes and metabolites in YH_3 suggested their potential involvement in secondary metabolism of microalgae after sewage treatment. When citrate and pyruvate were utilized to strengthen ATP generation for high cAMP production, oxidative stress became more severe in cells resulting in lower cell viability [38]. Thus, in this case, microalgae can induce the downregulation of succinic acid, a metabolite in propanoate metabolism pathways, to offset the influence of external environmental changes on microalgae and maintain normal metabolism.

## 5. Conclusions

*Desmodesmus* sp. from high altitude area are beneficial in wastewater treatment. In the process of studying the mechanism of sewage treatment, we found that the photosynthesis, oxidative phosphorylation, and propanoate metabolism signaling pathways were the most significantly enriched pathways in response to sewage treatment as shown by the combined transcriptome and metabolome analysis.

In addition, we found that *Desmodesmus* sp. response to sewage treatment could also induce numerous changes in primary metabolism, such as carbohydrate, fatty acid, and amino acid metabolism as compared to the control.

Overall, the results described here should improve fundamental knowledge of molecular responses to *Desmodesmus* sp. in sewage treatment and contribute to the design of strategies in microalgal response to sewage treatment.

## Supplementary Materials

The supplementary materials are available online. Fig. S1: GO annotation classification statistics chart; Fig. S2: Statistical map of KEGG metabolic pathway classification; Table S1: All metabolites detected by metabolome in positive polarity mode. Table S2: All metabolites detected by metabolome in negative polarity mode. Table S3: The differentially expressed genes and differentially accumulated metabolites are simultaneously mapped to the KEGG pathway diagram in positive polarity mode. Table S4: The differentially expressed genes and differentially accumulated metabolites are simultaneously mapped to the KEGG pathway diagram in negative polarity mode. Table S5: The differentially expressed genes and differentially accumulated metabolites are simultaneously mapped to the KEGG pathway.

## Author Contributions

Conception and design of the study: W.J., Z.Q., B.D. Investigation: W.J., Z.Q., C.J., Z.J., L.J., W.Y. Data analysis: W.J., Z.Q., Z.J., W.Y. Manuscript writing: W.J., B.D. All authors read and approved the final manuscript.

## Acknowledgments

We thank Novogene Bioinformatics (Beijing) Technology Co. Ltd. for the sample processing, extraction, and metabolites detection for metabolome analysis following their standard procedures.

